# Short-term monocular deprivation in adult humans: a meta-analysis and new perspectives

**DOI:** 10.1101/2025.02.20.639298

**Authors:** Cecilia Steinwurzel, Giacomo Pennella, Paola Binda

## Abstract

Starting from the early 2010s [1, 2], several studies have shown that a short period of monocular deprivation in adult volunteers transiently shifts ocular dominance in favor of the deprived eye. We compiled a meta-analysis of 73 such studies that measured the effects of monocular deprivation and related manipulations, using a diverse set of techniques [1-73]. The ocular dominance shift elicited by monocular deprivation was comparable across studies where deprivation was achieved with an opaque or translucent patch and irrespectively of whether the dominant or non-dominant eye was deprived. Effects were larger and longer lasting after longer periods of deprivation. Qualitatively similar effects were produced by monocular manipulations that did not reduce the strength of the stimulus: filtering or distorting the image in one eye, suppressing it from awareness or merely making it task irrelevant. We discuss the available evidence in the light of current models and a new perspective inspired by predictive coding.

## Introduction

Monocular deprivation is a classic paradigm used to study neuroplasticity. During development, blocking vision in one eye for several days triggers a permanent shift of ocular dominance in favor of the non-deprived eye [74-76], i.e. amblyopia. The molecular mechanisms of the effect likely involve both Hebbian plasticity (mainly, long-term depression of the deprived eye connections) and homeostatic plasticity [77-80]. The same monocular deprivation performed in adult animals generally fails to affect ocular dominance in any appreciable way [80]. However, a shorter deprivation (minutes or hours) in adult humans does promote a transient shift of ocular dominance in favor of the deprived eye, i.e. the opposite of what happens during development. Here we meta-analyzed a large set of studies that documented this effect. We started from the study by Lunghi et al. in 2011 [1], followed by Zhou et al. in 2013 [2], which opened a line of research demonstrating the residual plasticity of the adult visual system. We introduce this by reviewing a few antecedents of this line of work.

To our knowledge, the earliest investigations on the effects of monocular deprivation on human adults’ vision were carried out by Zubek & Bross in the 1970s [81-83]. They showed that monocular occlusion produced a biphasic modulation of temporal resolution in the non-deprived eye, which was briefly impaired in the first few hours and then enhanced for several days. This effect was specific for monocular deprivations achieved with a light-tight patch and it was selective for the non-deprived eye, with no change in the deprived eye, making it very different from the effects reported by the 2011 [1] and later publications.

Another relevant set of studies was performed by Xu et al. [84, 85], who tested the effect of repeatedly removing stimulation from one eye (1h a day for 10 days) and compared it with the effect of merely cueing attention to stimuli in one eye. Both manipulations produced a transient shift of ocular dominance in favor of the unstimulated eye (like in [1]) or the unattended eye, observed immediately after the 1h manipulation. However, there was also a persistent effect that built up over the 10 days, selectively observed for the attentional manipulation and shifting ocular dominance in the opposite direction: in favor of the cued eye. This is strong evidence that the adult visual system retains the ability to recalibrate in response to altered visual statistics, which was also supported by the application of binocular image distortions. For example, it is common experience that wearing new prescription glasses comes with a measure of discomfort, which subsides within days. Adams et al. [86] provided a quantification of this experience, demonstrating a gradual adjustment of 3D slant perception. Neitz et al. [87] demonstrated that biasing the spectrum of everyday light exposure shifts our color perception. Both these studies indicate that fundamental properties of our visual experience, like space and color, can change depending on experience throughout our adult life. Can ocular dominance adjust as flexibly? Or is the transient ocular dominance shift elicited by short-term monocular deprivation a qualitatively different phenomenon? Our meta-analysis is a first step towards addressing these questions. Although amblyopia treatment is a major motivation behind this line of research, we only included results from normally sighted adults; for a clinical perspective on this literature, the reader is referred to two recent reviews [88, 89].

## Methods

### Search protocol and exclusion criteria

We followed the PRISMA statement [90] and checklist guidelines. On November 28^th^, 2024, we conducted a PubMed search for the keywords ‘Monocular Deprivation’ or ‘Monocular Patching’ or ‘Ocular dominance plasticity’, excluding publications before 2011 and all reviews/editorials. This gave a total of 673 studies, from which we manually excluded all publications involving animal models or patients and focused on the remaining 72 studies of short-term monocular deprivation in normally sighted adult humans. To these, we manually added 1 publication [10] that was not identified by the initial search but was cited in the selected studies, yielding a total of 73 studies. Of these, 54 studies contributed to at least one dataset for our meta-analysis. Datasets were excluded when: 1) the experiment did not measure ocular dominance, 2) the reported figures/statistics/supplementary data did not allow for converting the ocular dominance shift into an effect-size estimate (defined in the next section), 3) the dataset included less than 4 participants, 4) the dataset was described as a re-analysis of a previous study (i.e. we excluded duplicates), and 5) the monocular deprivation was performed in non-standard conditions (pharmacological manipulations, brain stimulation, physical activity, video-game playing, marked alterations of the visual experience like staring at a blank surface, binocular deprivations, and others). Supplementary Table 1 lists all datasets and motivates exclusions according to these five categories.

### Effect-size computation

We defined effect sizes based on the “deprivation index”, which quantifies the change of ocular dominance after the monocular manipulation. For many experiments, we used the deprivation index reported in the source publication; this was either computed as a post-to pre-deprivation difference (e.g., for dominance proportions in binocular rivalry) or as a ratio (e.g., for mean phase durations in the same task). We applied a logarithmic transformation to ratio values to ensure that, in both cases, a null effect is represented by a deprivation index of 0 and a positive deprivation index implies a shift of ocular dominance in favor of the manipulated eye. Some experiments did not report a deprivation index but reported ocular dominance values before and after deprivation. Ocular dominance was either measured as the proportion of time during which one eye dominated perception (e.g., in binocular rivalry), as the bias resulting from binocular combination of partly incongruent visual stimuli (e.g., phase shift), or as the log-ratio between values describing vision in each eye. In these cases, we computed the deprivation index by taking the pairwise difference of the post-deprivation to pre-deprivation values. Finally, some experiments reported values indexing vision in the two eyes separately (e.g., mean phase duration in binocular rivalry or monocular contrast sensitivity), raw or as ratios to their pre-deprivation baseline; in this case, we computed the deprivation index as the ratio of the change in the two eyes (post/pre ratio in the deprived eye, divided by the same for the non-deprived eye) and log-transformed the results.

Despite these transformations, results from different experiments remained heterogeneous in their units of measurement: some did not have any (all ratios), others expressed the deprivation index as a proportion (dominance in binocular rivalry) or in degrees of visual angle (phase-shift in binocular combination). We therefore performed one last transformation, expressing the average deprivation index in each experiment in units of its standard deviation across the participants’ sample – like in Cohen’s *d* [91]. This allowed us to compare the efficacy of different monocular manipulations. To extract effect-sizes, we used one of three sources: (1) individual participants’ values, as shown in figures or reported in online repositories; (2) paired t-tests comparing ocular dominance before and after the monocular manipulation (we divided the t-statistics by the square root of the sample size minus one, transforming it into our effect size estimate); (3) bar-plots reporting mean and standard errors of the deprivation index (we transformed the standard errors into standard deviations multiplying them by the square root of the sample size minus one). When none of these options was available, the experiment was excluded from our main analyses (see Supplementary Table 1). In several publications, the effects of monocular manipulations were evaluated at multiple time-points. For the above analyses, we used the first measurement that usually coincides with the peak effect – but note that studies differ markedly in the duration of the time-window over which this first measurement was averaged, from less than 3 minutes [2] to more than 10 minutes [67]. In addition, we pooled across experiments reporting multiple time-points to examine the decay rate of the effect, considering only data from the two most common techniques (binocular rivalry and binocular phase combination) with black-and-white stimuli (to minimize the impact of color, reported in [5]). These reported effects in one of three ways: (i) with deprivation index values computed as post to pre-deprivation differences and reported on a linear scale, e.g. [2]; (ii) with ocular dominance values after deprivation, expressed as ratios between eyes and reported on a linear scale, e.g. [1]; (iii) with deprivation index values computed as post to pre-deprivation ratios and expressed in logarithmic units or as decibels, e.g. [28]. We expressed all values on a linear scale and ensured that the expected baseline is 0. This required making the following transformations: (i) no changes; (ii) subtracting 1 to bring the expected baseline to 0; (iii) transforming logarithmic values into linear ones (after dividing by 20 values expressed in decibels) and then subtracting 1 to bring the expected baseline to 0. This produced a series of plots where the effect decays exponentially with the delay from the end of the manipulation. We represented these in a log-log space, fitting each curve with a linear function, and used its slope to estimate the inverse of decay rate (for visualization purposes, the curves in Figure 2B are also normalized to the peak effect).

### Software and statistical approach

When extracting data from published figures, we used the software WebPlotDigitizer (https://automeris.io/WebPlotDigitizer/). Statistics were performed in Matlab [92] and JASP [93]. JASP was used to estimate a Linear Mixed Model on effect-sizes for different techniques, after accounting for the influence of the variable “delay of test from patch removal”, which was treated as a random factor. Pearson’s correlation coefficients we estimated using the “robust correlation toolbox” [94], which identified outliers using the “boxplot” option.

## Results for monocular deprivation studies

The selected 73 studies contributed 251 datasets from independent experiments. The majority of these (N=192) tested the effects of a brief period of monocular deprivation achieved through the application of a patch that completely (light-tight patch) or partially (translucent patch) occluded vision in one eye, or by nulling contrast in one eye under dichoptic viewing conditions [12, 21, 34], which we consider conceptually equivalent to the application of a translucent patch. From these, 111 experiments measured the effect of monocular deprivation (without any intervening manipulations) on ocular dominance (rather than other indices) and reported sufficient data for us to compute effect sizes, which we report in Figure 1.

**Figure 1.**
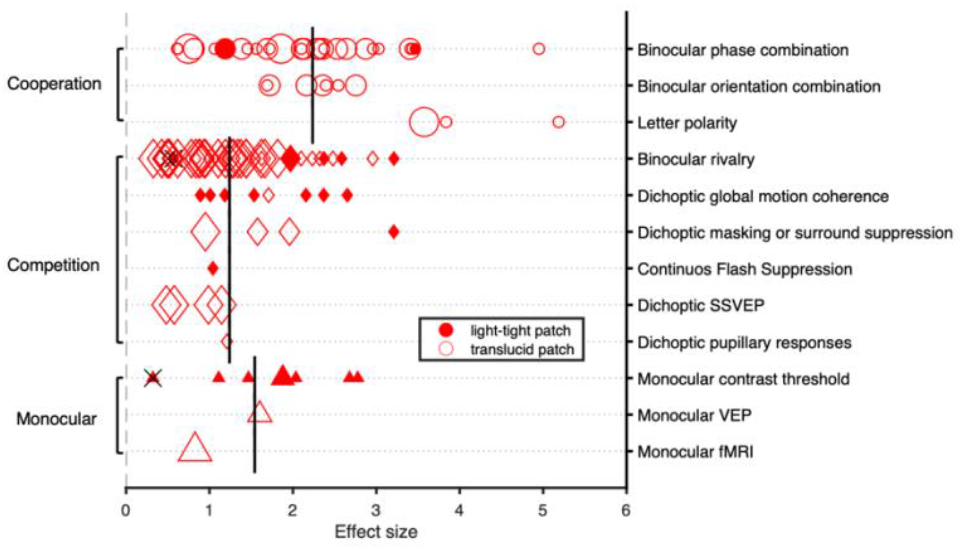
Effects of monocular deprivation on ocular dominance (x-axis) measured with different techniques (y-axis). We grouped these into three categories (marker shape, used in all other figures) for which the average effect is given by the vertical lines. Marker color (filled or empty) indicates the type of monocular patch used (light-tight or translucent). Marker size represents sample size: small (<10 participants), medium (11–15 participants), or large (>16 participants). Crossed-out symbols represent non-significant effects

### Ocular dominance studies

Ocular dominance, i.e. the relative contribution of the two eyes to visual perception, may be estimated with a diverse set of techniques (see Supplementary Methods). We expressed the effects of monocular deprivation as the average ocular dominance shift in units of standard deviation – as conceptualized in the Cohen’s *d* index [91]. Where necessary, source data were transformed to ensure that positive effects represent a shift of ocular dominance in favor of the deprived eye and that the expected value under the null hypothesis that deprivation had no effect is 0. Pooling across experiments, the grand-average effect is large: 1.7 units of standard deviation, predicting that >95% of individuals in the population will display an ocular dominance shift in favor of the deprived eye.

Experiments using very different techniques (y-axis of Figure 1) yielded qualitatively similar results. We grouped techniques into three categories: stimulating each eye separately (monocular), stimulating the two eyes with incongruent images that do not fuse (binocular competition) or fuse into a unitary percept (binocular cooperation). Vertical lines report the average effect size in each category, showing comparable effects for monocular and binocular competition tests (t(68) = 0.90, p = 0.373) but larger effects for the binocular cooperation tests (binocular cooperation vs. monocular: t(48) = 1.72, p = 0.092; binocular cooperation vs. binocular competition: t(100) = 5.01, p<0.001). However, this difference was completely accounted for by a confounding factor, test durations. While most binocular cooperation experiments reported the ocular dominance shift in the first 3 minutes after patch removal, many binocular competition experiments averaged over >10 minutes. After accounting for the effect of test-duration (and excluding the 5 experiments that did not explicitly report this information), binocular cooperation and competitions effects were non-significantly different (linear mixed model with delay as random effect, Satterthwaite approximation: F(1,1.82) = 0.42, p = 0.586). These results indicate that the effect of monocular deprivation is robust and easily measured with both binocular and monocular techniques, with no reliable difference among these.

We also found no reliable difference between the effects of applying a light-tight (N=23) or a translucent patch (or contrast-nulling in dichoptic viewing, N=88; two-sample t-test: t(109) = 0.86, p = 0.391), in line with qualitative comparisons [2, 45]. This implies that preventing pattern vision in one eye is sufficient to shift ocular dominance, while eliminating all visual input (luminance) is not relevant.

A long-standing assumption is that depriving the dominant eye is more effective than depriving the non-dominant one, as originally reported by Lunghi et al. [1] (for 8 participants). Probably as consequence, we only found N=17 experiments depriving the non-dominant eye, while N=84 deprived the dominant eye (the remaining N=10 did not impose or report a systematic association between the deprived and dominant eye). However, a statistical comparison of the effect sizes revealed no reliable difference between depriving the dominant or non-dominant eye (t(99) = 1.03, p = 0.305). We acknowledge that this across-study approach has multiple limitations, including the unequal sample sizes and the heterogeneity of criteria used to identify the dominant eye (for example, some used pre-deprivation binocular rivalry or phase-combination results, others used separate tests like the Porta test). Further within-subject studies are warranted.

In Figure 1 there are only 3 datasets with a non-significant ocular dominance shift [25, 38, 56]. These null findings are counterbalanced by several positive results obtained with similar techniques and deprivation approaches, suggesting that they may be related to experimental noise.

#### The dose-response curve of the monocular deprivation effect and its decay-rate

Is the ocular dominance shift elicited by monocular deprivation dependent on deprivation duration?

Eight studies tested more than one deprivation duration [5, 24, 29, 43, 45, 52, 56, 63]; four of these provided strong evidence for a larger effect at longer durations [29, 52, 63], according to a saturation function ceiling around 5h [43]. The other four studies gave more uncertain results: in two cases, effect sizes were similar across tested durations [24, 45]; in another, there was a positive correlation between duration and effect size but only when ocular dominance was measured with binocular rivalry, not with a letter polarity test [56]; finally, one study reported that the effect at 2.5h was larger than at 30min but also more variable, resulting in a paradoxically smaller effect size [5]. To explore this question further, we leveraged the variance in deprivation duration across studies (Figure 2A) and found a strong positive correlation between deprivation duration and effect size (Pearson’s r calculated after the automatic exclusion of two outliers, marked by gray symbols). Thus, evidence from both within- and across-study analyses consistently indicate that longer monocular deprivations induce stronger ocular dominance changes.

**Figure 2.**
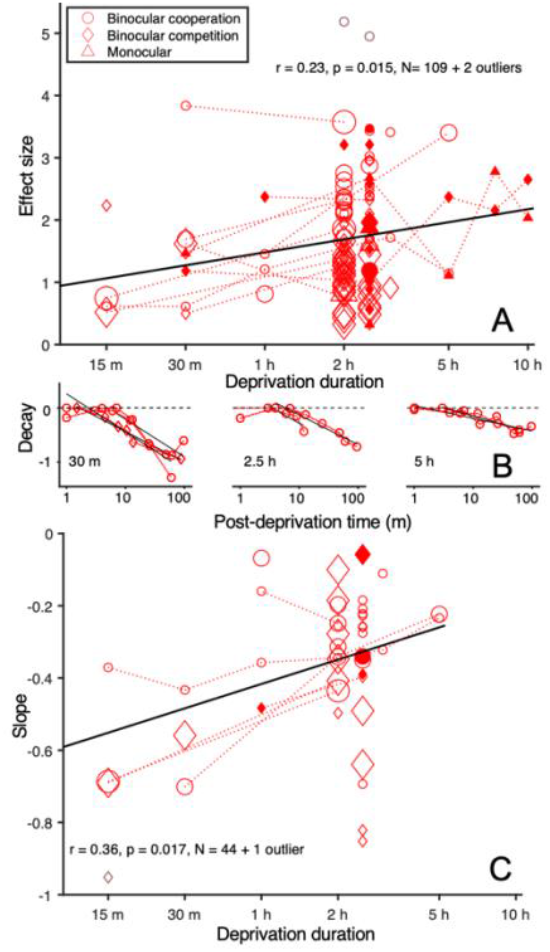
**A)** Relationship between deprivation duration and effect size. Marker shape, color and size follow the same conventions as in Figure 1; connected data-points are observations contributed by the same study. The black line shows the best fit linear function after automatic exclusion of the two data-points shown in gray. **(B)** Decay dynamics of the effect for studies that performed multiple post-deprivation measurements. Each subplot represents data from one deprivation duration (three example durations, specified by the text insets). Continuous lines show the best fit linear function through the double-logarithmic transformation of the normalized decay curve. When the peak effect was not recorded at the first time-point (i.e. immediately upon patch removal), the linear fit started from the peak and ignored earlier timepoints. The goodness of fit was adequate, averaging 0.84+/-0.02% variance explained (mean and standard error across all analyzed experiments). **(C)** Slope of the decay curves plotted against the deprivation duration. Same conventions as in Figure 1; data-points contributed by the same study and obtained with the same technique (binocular rivalry or binocular combination) are connected.

A related question is whether longer deprivations promote longer-lasting effects. To our knowledge, no study addressed this question, which we tackled by considering the subset of experiments for which the decay rate of the deprivation effect could be estimated (we restricted this analysis to the binocular rivalry and binocular combination techniques). Source data were transformed to ensure that deprivation effects were measured with homogenous units (see Methods), and we studied the slope of the function relating effect-sizes to delay of testing, both represented in a log-log space. As shown in Figure 2B, longer deprivation durations were systematically associated with slower decay rates, i.e. less negative slopes. This is quantified in Figure 2C, where the slopes are positively correlated with durations, implying that longer deprivations induced longer lasting effects.

Note that the analysis in Figure 2C is conceptually independent from that in Figure 2A, given the evidence that the decay rate can vary independently of the effect size. For example, using colored binocular rivalry stimuli, the duration of the effect can be extended (compared with achromatic gratings, expressed in the units of Figure 2C, slopes went from -0.54 to -0.09) without altering the peak effect [5]. On the other hand, physical activity [10] can increase the peak effect without altering its decay (−0.28 and -0.22 in the units of Figure 2C). Moreover, two studies asked whether the monocular deprivation effect can be “kept in storage” by interrupting all visual stimulation at the end of deprivation and found that dark exposure can prolong the effect for about one hour [61] (not two hours [51]); however, a much longer retention interval is possible if participants go to sleep at the end of the deprivation [51].

These results suggest that the decay of the deprivation effect is governed by multiple factors. Since the decay is not constant (it is delayed with longer deprivations and by going to sleep [51]), restoration of the normal visual input at the end of deprivation cannot be the only governing factor; of course, this remains an important factor, given that total darkness slows down the recovery of normal ocular dominance [61].

A final related point is whether the effects can be integrated across separate monocular deprivation sessions. Considering the typical decay rate, one would estimate that the effect of a typical 2-hour deprivation (slope about -0.3) would be halved in the first hour, reduced to 1/4 in 2 hours, 1/8 in 3 hours, 1/16 in 4 hours, etc. Assuming linear behavior, this predicts that monocular deprivations separated by few minutes should integrate their effects, as shown in the three studies that briefly interrupted deprivations to track their effect over time and reported a gradual increase of the effect-size [34, 43, 45]. It also predicts that no residual effect would be measurable 24 hours after the end of a monocular deprivation [12] and that the effects of monocular deprivations performed 24 hours apart should be indistinguishable [28, 73]. Note, however, that multiple daily sessions of amblyopic-eye deprivation promote a gradual recovery of its vision [95-97]. This suggests that monocular deprivation has physiological consequences that go beyond the transient shifts of ocular dominance and may outlast them, with important clinical implications.

#### Factors interacting with monocular deprivation

Figures 1-2 selectively considered experiments performed under very homogeneous conditions, which we clarify here by opposition with other studies that purposedly altered such conditions to test the impact of ***<u>within-subject</u> <u>factors</u>***. While most studies ensured that visual input to the non-deprived eye had similar statistics as in normal viewing, one study dramatically **reduced visual information in the non-deprived eye** by letting participants watch a blank curtain for the entire deprivation duration [45]. In these conditions, the non-deprived eye received little more information than the deprived eye and, predictably, the deprivation produced no reliable ocular dominance shift. Coherently, the deprivation effect was reduced when ocular dominance was measured with stimuli that were unattended during the deprivation, although presented to the non-deprived eye, they were actively ignored [71].

While most studies achieved monocular deprivation by covering one eye with a patch, behind which the eye remained open, one study assessed the impact of **keeping the deprived eye shut** [57] and reported a reduction of the deprivation effect. Shutting the deprived eye achieves a stronger deprivation, yet it produces a smaller effect.

Several studies manipulated the **activity performed during deprivation**, with mixed results. Engaging in moderate physical activity during the deprivation enhanced the effect in one study [10] but not in others [20, 26, 44, 46, 47]. Engaging in an orthogonal task (motor-sequence learning or working memory) reduced the deprivation effect in one study [64]; however, no modulation of the effect was observed when manipulating participants’ engagement in a visuomotor task (playing action videogames, versus passively watching the game [31]). This heterogeneity suggests that a latent variable related to the experimental manipulations may have been incompletely controlled – for example, the visual consequences of these activities (e.g. amount of gaze movement during the activities) may require more detailed characterization.

Another set of studies manipulated **neural excitability during deprivation**. There was no effect of applying non-invasive stimulation of the occipital cortex during deprivation [47, 56]. However, the deprivation effect was enhanced when (i) participants stayed in total darkness for one hour before the deprivation [61], (ii) the deprivation was performed in the morning vs. the late evening [51] and (iii) participants had a standardized breakfast before the deprivation vs. maintained the overnight fasting [55], all conditions associated with a modulation of cortical excitability (enhanced by dark exposure [98, 99, 100], increased in the morning vs. evening [101], and after a meal vs. fasting [102]), suggesting that enhanced cortical excitability facilitates the ocular dominance shift induced by deprivation. In contrast, the deprivation effect was reduced by administration of donepezil, a cholinesterase inhibitor [29], suggesting that increasing acetylcholine levels prevents ocular dominance from shifting in response to deprivation.

Few studies considered ***<u>between-subject</u> <u>covariates</u>*** and generally failed to detect systematic inter-individual differences in monocular deprivation effects, with one exception: **body mass index**, which is negatively correlated with effect size [27, 39]. Deprivation effects did not vary systematically with **age** [41, 62], as the ocular dominance shift following monocular deprivation was comparable in adolescents, young adults and elderly participants – but note that a stronger effect in the elderly participants could be revealed by onset rivalry [41]. No study reported association between **sex or gender** and monocular deprivation effects. One study found that the deprivation effect correlated with baseline binocular vision [37] but this correlation was not significant in other studies [45, 62]. These null findings do not exclude the existence of associations between monocular deprivation effects and important dimensions of inter-individual variability; however, they indicate that, if they exist, these associations are not strong enough to be revealed with the small sample sizes that are typical of this line of research.

### Other techniques and indices

So far, we only considered studies that tested the effects of monocular deprivation on ocular dominance; here we describe experiments that considered other aspects of perception or resting state physiology.

The same binocular tasks that measure ocular dominance also give indices of **binocular fusion** (or lack thereof, i.e. diplopia). Of the 20 experiments that reported such indices, 16 found no change [1, 5, 12, 17, 25-27, 37, 44, 45, 47, 55, 56, 62, 63, 68]; three found increased probability of mixed percepts after monocular deprivation [30, 41, 71] and one reported decreased mixed percepts [34]. These statistics suggest that binocular fusion is not strongly affected by deprivation. A separate set of tasks measured **temporal vision** and found that resolution is unaffected by monocular deprivation [32] (in partial contrast with the seminal work by Zubek [81, 82]); however, there was a slight delay (by few milliseconds) of the deprived eye relative to the non-deprived eye revealed through the Pulfrich effect [42, 53]). Three studies asked whether, besides changing monocular responses and binocular vision, monocular deprivation also affects the **integration of visual and non-visual signals**. Different results were obtained for different integration tasks. On the one hand, the temporal binding window for visual and auditory events was enlarged for the deprived eye, suggesting enhanced audio-visual integration [35] (although this was insufficient to affect the sound-flash illusion [58]); on the other hand, visuo-haptic integration during rivalry was impaired for the deprived eye, suggesting that monocular deprivation reduces our ability to integrate information across senses [16].

Finally, five studies measured how monocular deprivation affects **resting state** physiology. Occipital alpha rhythms are clearly affected; induced alpha during stimulation of deprived eye was reduced [58] and resting alpha during pauses between deprived eye stimulations enhanced [8]. Combined with the observation that GABA concentrations in primary visual cortex are reduced after monocular deprivation [9], these results provide strong indications that monocular deprivation is linked with enhanced cortical excitability (see also above). One apparent contradiction is with an early TMS study, which reported decreased probability of phosphene generation during application of a monocular light-tight patch [3]; however, it is unclear whether the conditions of this study are entirely comparable with the rest of the meta-analyzed studies.

Pupillometry was also employed to measure changes of the resting-state physiology across monocular deprivation. This study revealed that a slow rhythm (<1 Hz), termed “hippus” and related to the balance of autonomous nervous signals, is reliably enhanced following monocular deprivation [13]. One final study investigated the impact of monocular deprivation on perceptual learning. Shibata et al. [4] applied monocular deprivation continuously for three days and reported an acceleration of learning for the non-deprived eye – suggesting that monocular deprivation may promote our ability to learn from the environment.

In summary, studies that did not look at ocular dominance applied a heterogenous set of techniques and produced partially conflicting results. However, they have the important merit of showing the complexity of monocular deprivation effects, which likely extend beyond a gain modulation of visually evoked responses.

## Results for other monocular manipulations

While most studies tested the effects of blocking vision in one eye, a growing number of experiments are exploring more subtle manipulations and surprisingly report essentially the same effect: a shift of ocular dominance in favor of the manipulated eye. Of the 59 available datasets, 37 allowed us to compute an ocular dominance effect-size and are reported in Figure 3.

**Figure 3.**
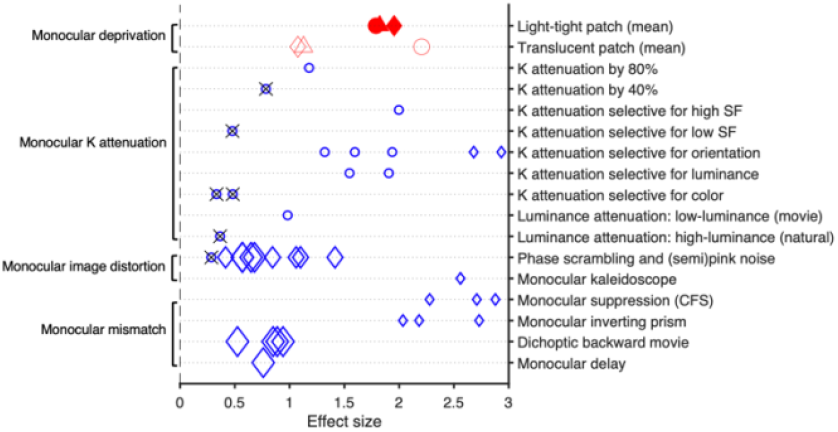
Distribution of the effects of monocular manipulations on ocular dominance, obtained using non-standard patching deprivation methods. The X-axis represents effect size (Cohen’s d). Each data point corresponds to a study employing alternative deprivation techniques, as listed on the right Y-axis. Symbol size and shape use the same conventions as in Figure 1. Symbol size for mean values of translucent and light-tight patching are standard and not reflecting the effective sample size. K: contrast. SF: Spatial Frequencies. CFS: Continuous Flash Suppression.

One manipulation that is conceptually close to monocular patching consists of **partially attenuating contrast in one eye** (in a dichoptic stimulation set-up). The attenuation needed to be strong (80%, not 40%) to elicit a reliable ocular dominance shift [7]. Instead, swapping the contrast attenuation between eyes (at 7Hz, [50]) did not affect ocular dominance, indicating that interfering with binocular fusion is not sufficient to elicit the shift. Selective attenuation of luminance contrast (leaving chromatic contrast unaffected) was sufficient to elicit a strong ocular dominance shift, while the opposite manipulation (attenuating chromatic contrast, leaving luminance contrast unaffected) was ineffective [21]. Attenuating contrast at one range of orientations (e.g., vertical or horizontal) elicited a strong ocular dominance shift, unselective for the orientations used for testing [18]. A strong effect was also achieved attenuating contrast of the higher spatial frequency components, again unselective for the frequencies used for testing; however, no such effect was elicited by attenuating the lower spatial frequency components [7]. This observation is consistent with the effects of a monocular neutral density filter, which elicited a strong ocular dominance shift only when participants viewed artificial (low luminance) stimuli, not in natural viewing conditions [19]. These results collectively indicate that monocular deprivation primarily acts through reduced visibility of the high-frequency components of the image. Higher spatial frequencies are associated with lower contrast sensitivity, implying that they are most impacted by contrast attenuation. Moreover, acuity is lower for iso-luminant stimuli defined by chromatic contrast, explaining why a strong ocular dominance shift is achieved by depriving luminance contrast (leaving only iso-luminant chromatic modulations that do not efficiently carry the higher spatial frequencies). When average luminance decreases through a neutral density filter, the contrast sensitivity function shifts to the left, selectively attenuating sensitivity for the higher spatial frequencies (but this effect is negligible in natural light, where luminance is so strong that the filter has no appreciable impact on acuity).

Without attenuating contrast, an ocular dominance shift could be induced by **distorting the image in one eye**, by phase-scrambling the image presented to one eye [7], by substituting it with pink noise [12, 34, 67], or by applying a monocular kaleidoscope [25]. And even without altering the monocular image in any way, the ocular dominance shift could be induced by **making it invisible** [15, 33] through continuous flash suppression (note that this effect was already present in the 2010 report by Ooi et al. [84]).

A final set of studies merely **mismatched the monocular image with its multimodal context**. This was achieved in one of three ways: (i) with a monocular inverting prism [45]; (ii) by reversing the temporal order of movie-frames in one eye [65, 69, 70]; (iii) by delaying one eye by 1/3 of a second to make it useless for visuomotor coordination [66]. In all three cases, ocular dominance shifted in favor of the manipulated eye (note that these manipulations only lasted 1h, about half the typical duration of monocular deprivation, complicating the comparison of effect size across manipulations). Control experiments assigned a key role to behavioral relevance for the generation of the ocular dominance shifts; for example, monocular delay was ineffective when participants passively viewed the scene rather than engaging in a visuomotor task [66]. Moreover, the effect of temporal inversion was depleted by non-invasive inhibition of the Frontal Eye Fields [70].

It is interesting to consider the few examples where multiple techniques were used to test the same or similar manipulations. This was the case for monocular image distortion, where the ocular dominance shift was selectively observed with binocular competition techniques, not with binocular cooperation [7, 12]. Based on a multi-stage model of binocular processing [103], these authors suggested that monocular distortion could impact ocular dominance at a later stage than monocular deprivation, sparing monocular and binocular cooperation stages. However, this model is inconsistent with data from the monocular kaleidoscope manipulation (another type of monocular distortion), which produced similar effects on both binocular competition and monocular contrast sensitivity [25].

It is also informative to compare studies testing the effects of different manipulations with the same technique. Binocular rivalry and steady-state visual evoked EEG responses (elicited by incompatible stimuli in the two eyes) were tested for two manipulations: monocular deprivation (nulling contrast in one eye [67]) and monocular mismatching (dichoptic backward movie [70]). Both manipulations produced similar effects with both techniques. However, measuring evoked potentials with compatible binocular stimuli only revealed an effect of monocular deprivation [34] and monocular distortion (presenting pink noise in one eye [34]), but no effect of the dichoptic backward movie manipulation [65]. Some of these studies additionally analyzed intermodulation frequency responses, which estimate the strength of inter-ocular competition. This was reduced by the dichoptic backward movie manipulation [70], but unaffected by monocular deprivation [67].

## Discussion

In the nearly 15 years of short-term monocular manipulation studies, several models have been proposed. One discarded hypothesis is that **adaptation to low contrast** can account for all effects. A recent review listed many qualitative differences between contrast (de-)adaptation and monocular deprivation [104]. For example, the sharper vision experienced after adaptation to low contrast [105] or to blur [106-110] transfer across eyes and tend to be tuned for orientation [111, 112], the exact opposite of the effects of monocular manipulations [15, 18]. Besides, adaptation cannot explain the effects of monocular manipulations that do not affect monocular contrast (e.g., [45, 65, 66]).

The most common hypothesis describes the effects of monocular deprivation as a form of **homeostatic plasticity**. This hypothesis can be specified by assuming that monocular deprivation promotes a gain-change in monocular processing, enhancing gain for the deprived eye and/or decreasing gain for the non-deprived eye. In principle, the change could either impact contrast-gain (shifting the contrast threshold without affecting the maximum response) or response-gain (scaling response amplitude with little impact on contrast thresholds). Experiments showing changes of monocular contrast sensitivity [1, 2, 8, 11, 21, 23, 25, 38, 43, 49, 57, 58, 60] land some support to the contrast-gain model. One study [60] applied the “Perceptual Template Model” and found that enhanced sensitivity in the deprived eye is best explained by reduced additive internal noise, which is conceptually close to a contrast-gain enhancement. In both cases, the effect is compatible with a change of the signal-to-noise ratio in early visual processing, predicting that short-term monocular deprivation should modulate the inhibitory tone in the visual cortex, in line with MR spectroscopy findings [9]. Based on these, many have interpreted the effects of monocular deprivation as resulting from reduced GABAergic inhibition in early visual processing [41, 55, 62, 66]. While this can directly account for the enhanced deprived-eye responses, it is not immediately related to the suppression of the non-deprived eye, which would presumably result from further processing such as reduced interocular inhibition. In line with this, some proposals assume that the effects of monocular deprivation only emerge at the level of interocular combination, without directly affecting monocular processing. One specific formulation [65, 67, 70] is based on a binocular integration scheme featuring “opponency neurons” that mediate inter-ocular inhibition [103]. Monocular deprivation would induce the selective adaptation of opponency neurons activated by the non-deprived eye, removing inhibition over the deprived eye and accounting for its boost. While this model fits well the results from a complex monocular manipulation (dichoptic backward movie), it conflicts with some of the results from standard monocular deprivation – for example, deprivation fails to affect “intermodulation frequencies”, a key index of interocular inhibition [67]; moreover, as shown in Figure 1, the effects of monocular deprivation are readily measurable by monocular stimulation (whether this is the case for the more complex manipulations in Figure 3 is currently unknown). In general, the homeostatic plasticity accounts model the effects of monocular manipulations as local gain changes within the primary visual cortex V1, the last cortical stage where information from the two eyes is represented separately. Such local-V1 perspective does not easily account for several complex/high-level factors that have been linked with the ocular dominance shifts. These include the observation that multisensory integration fails for the deprived eye [16] and that monocular deprivation effects are modulated by motor and cognitive tasks [64]; these clearly suggest that signals from outside V1 participate in the effect, in line with evidence that monocular deprivation affects feed-back modulatory signals [58]. It is also interesting to explore the accounts that such local-V1 models could give of the effects of complex monocular manipulations, different from monocular deprivation. Song et al. [65] assumed that opponency neurons are modulated by attention, outlining a circuit-model where signals related to complex information, like the behavioral relevance of the image, originate outside V1 (e.g. in FEF, as suggested by a non-invasive brain stimulation results [69]), but eventually feed into V1 to affect the balance between the two eyes. This leaves open the crucial question how FEF would know which opponency neurons to target. Another possibility is that complex monocular manipulations modulate V1 activity in ways that mimic what happens during monocular deprivation. For example, when one eye is unattended or unnoticed (due to continuous flash suppression [15] or application of a spatiotemporal inversion/delay [45, 65, 66]), its cortical representation could be suppressed; at the end of the manipulation, this suppression could rebound into the observed boost of the manipulated eye (as suggested in [15], [45] and in the early work by Ooi et al. [84]). This model assumes that an internal gain-control regulation follows a very complex time-course, gradually shifting in one direction during the manipulation and suddenly swapping to the other direction at the end of the manipulation. A simpler model could make the opposite assumption: that V1 responses to stimuli outside attention or awareness are gradually enhanced during the manipulation. Although counterintuitive, this model is in line with the empirical observation that binocular rivalry tests performed during the manipulation suggested an *enhancement* of the unattended eye [45].

We take this opportunity to present a novel hypothesis that incorporates elements of the above models but starts from a radically different assumption: that **predictive coding** lays at the foundation of monocular manipulation effects. Predictive coding is a popular theory of neural function (specified in different models [113-117]) assuming that an important share of neural responses do not represent sensory inputs, but their deviations from predictions. These responses, for example in V1, will be enhanced when the input is inconsistent with predictions, suppressed otherwise. Over a lifetime of binocular experience, our visual system sets up a strong prediction that the two eyes provide congruent information. When one eye suddenly fails to match this prediction, it generates error signals. And if errors repeatedly occur for minutes or hours, the natural (Bayesian) consequence would be a downweighing of predictive signals [118] for the manipulated eye representation, explaining the observed enhancement. Like in all other models, the core effect is a gain-change in primary visual cortex, where the two eyes are represented separately, and the driving force is reduced inhibition. However, in this model inhibition implements the suppressing action of predictions (suppressing responses to predictable inputs) and its reduction implements the downweighing of predictive signals. Crucially, by assuming that the origin of the deprivation effect lays in modulatory signals from outside V1, this model readily accounts for the observed modulations of these connections [58]. We note that computational models [119] show how predictive coding systems can solve the difficult problem of correctly crediting mismatched predictions, so that errors generated at a high level can be routed to the V1 units representing one eye, not the other. This model accounts for the comparable effects of applying a light-tight or translucent patch, as predictions relate to the image content, not its luminance; it may also account for the preferential involvement of high spatial frequencies, as these are the main source of the information that we use for making sense of the visual scene, i.e., the main source of predictions. Within- and across-subject factors that modulate the efficacy of the manipulation could do so by altering our “predictive state”; for example, active engagement in a multisensory task could generate stronger predictions than passively watching a screen; in contrast, closing the deprived eye behind the patch could attenuate the mismatch, through the corollary discharge associated with lid-closure [120, 121]. This model could provide a unifying account for all monocular manipulations, both patching and more complex manipulations, suggesting that a single factor may be responsible for the manipulated eye enhancement: unpredictability. Deprived, filtered, distorted, inverted and delayed images are all unpredictable due to their incongruency with the rest of the sensorimotor input. This predicts that monocular mismatch and monocular deprivation should produce comparable and correlated effects; although there is some evidence for discrepancies in effect-size, the available data is not conclusive.

One final note concerns the relation between the short-term effects of monocular deprivation and long-term plasticity effects. The most important case is the persistent recovery of amblyopic eye acuity achieved by repeatedly patching the amblyopic eye in adults [95-97]. To account for this, some authors recall the close connection between homeostatic and Hebbian plasticity observed during development [88] – however, the details of this model have not been spelled out. Our predictive coding hypothesis could be extended to describe amblyopia as a case of inadequate predictions. The visual impairment in one eye during development could lead to excluding this eye from the error-prediction cycle, thereby cutting it off from perception. This implies that the predictive system could be responsible for the suppression of amblyopic eye responses when visual treatment is delayed after the critical period. It also implies that recovery in adulthood should be allowed by mismatching predictions for the amblyopic eye. As this may be achieved with a variety of techniques (including amblyopic-eye mismatching), our proposal may open new pathways for adult amblyopia treatments inspired by predictive coding.

