## Supplementary methods for "Short-term monocular deprivation in adult humans: a meta-analysis and new perspectives"

In what follows, citations follow the same numbering as in the main text.

### Binocular techniques

Binocular techniques afford a direct index of ocular dominance. This may be estimated in two ways: by presenting incoherent stimuli in the two eyes and testing how they are mutually suppressed (which we dub "binocular competition" tests) or by presenting relatively coherent stimuli in the two eyes and testing how they cooperate to support task performance (we refer to these as "binocular cooperation" tests).

Among the **binocular competition** techniques, **binocular rivalry** was used in the first study [1] and in many subsequent ones (please see Supplementary Table 1). In binocular rivalry, two incompatible images are presented to the two eyes and perception alternates between seeing through either eye or (more rarely) a combination of the two, the so-called mixed percepts. The effect of monocular deprivation (and other monocular manipulations) consists of an increased proportion of time spent seeing through the manipulated eye, and an increase of the manipulated eye mean phase duration. Min et al. [40] suggested that binocular rivalry has lower reliability than other binocular cooperation tasks and it is less effective for revealing the effects of monocular deprivation; however, inspection of Figure 1 suggests that the average effect size for binocular rivalry is above 1, more than four times larger than the effect size considered in [40]. A variation of the binocular rivalry technique is **onset rivalry**, measuring only the first phase of binocular rivalry, which is reliably biased towards the deprived eye after monocular deprivation [1, 41, 44, 62, 72] – possibly a larger effect than the ocular dominance shift computed from an extended period of rivalry [1, 41].

A related technique is **continuous flash suppression**, where one eye is presented with a dynamic, salient stimulus that steadily dominates over the less salient static image presented to the other eye; the contrast of the suppressed stimulus is increased until it "breaks suppression". Monocular deprivation decreases the contrast increment required for the deprived eye to break suppression [38], consistent with enhanced sensitivity in the deprived vs. non-deprived eye. The measures achieved by **dichoptic masking** are conceptually similar, as these techniques measure the sensitivity in one eye (e.g., the deprived eye) as a function of the presence and contrast of an interfering "mask" stimulus in the other eye. Masks generally elevate monocular detection thresholds; the effect of a mask in the non-deprived eye was reduced after monocular deprivation [22, 59]. In a similar experiment, the mask consisted of an annulus surrounding the target stimulus; target and mask were again presented in separate eyes (the deprived and non-deprived eye respectively), producing a **dichoptic surround suppression** effect, which was reduced after deprivation [46] and after a more complex monocular manipulation [33]. Finally, in the **dichoptic global motion coherence** task, the two eyes are presented with dots, some coherently moving in one direction (presented to the deprived eye), others moving randomly (presented to the non-deprived eye). The random motion in the non-deprived eye interferes with the detection of coherent motion; after monocular deprivation, this interference was reduced [2, 25, 43], once again consistent with enhanced dominance of the deprived eye over the non-deprived eye.

Studies using **dichoptic frequency tagging** with Electro- [11, 67, 70] or Magneto-EncephaloGraphy [14] reached similar conclusions. In this approach, the two eyes are shown with incongruent stimuli (e.g. orthogonal gratings) that flicker at different frequencies; the Fourier spectrum of the EEG or MEG signal is dominated by the first harmonics of both frequencies and their amplitudes measure the contribution of the two eyes to the steady-state visual evoked response. The ratio between responses in the two eyes was shifted in favor of the manipulated eye, both following monocular deprivation [11, 14, 67] and more complex manipulations [70]. The two studies that analyzed the modulation separately for each eye reported a selective enhancement of the deprived eye with no reliable modulation of the non-deprived eye [11, 14]. Acquafredda et al. [68] explored how **pupillometry** can serve as an objective physiological marker of ocular dominance plasticity by measuring pupil diameter during binocular rivalry before and after short-term monocular deprivation; they found an increase in the amplitude of these pupil-size modulations after deprivation – with no change in absolute pupil diameter, suggesting that the effects were not due to general retinal adaptation or changes in luminance sensitivity.

Among the **cooperation** techniques, **binocular phase combination** was the most common (please see Supplementary Table 1). The two eyes are shown with similar, though not identical stimuli, e.g. gratings with slightly different orientation or phase; the task is to report the phase or orientation of the single percept resulting from the fusion of the two images. Monocular deprivation increased the weight of the deprived eye stimulus in the fused percept. Like for binocular rivalry, stimuli can either be chromatic or achromatic. While most studies used achromatic stimuli, Zhou et al. [21] directly compared the effects measured with a/chromatic stimuli and found them indistinguishable; the same comparison with binocular rivalry revealed a difference in the dynamics of the effect [5], suggesting that the two techniques might measure different aspects of the monocular deprivation effect [12, 63]. A related implementation is the dichoptic tilted edges task [17], where the two eyes are presented with blurred edges of varying stimulus disparity and contrast. Participants report whether they perceive a single flat edge (indicating fusion), a single tilted edge (indicating dominance of one of the two monocular images and suppression of the other), or two tilted edges (indicating diplopia). While the probability of fusion and diplopia were unaffected by deprivation, the deprived eye image tended to dominate perception, suppressing the non-deprived eye. An alternative approach is the **binocular contrast matching** task [2], where the two eyes are presented with gratings of identical phase and orientation, but different contrast. The perceived contrast of the fused percept is intermediate between the two monocular images, as may be appreciated by matching it with a monocular grating of variable contrast presented in the deprived eye. A simpler variant of this task requires participants to match the contrast of two monocular stimuli presented at abutting locations [1]. In both cases, monocular deprivation shifted contrast matches indicating that the deprived eye required less contrast to match the stimulus in the non-deprived eye [1, 2].

Similarly, in the **dichoptic letter polarity** test [47, 56] each eye saw two letters, one white the other black and the contrast polarity of the letters is swapped across eyes. Participants reported which letter appeared brighter, indicating dominance of the eye that was presented with the white version of that letter. Monocular deprivation increased the proportion of trials where the letter presented in white to the deprived eye was chosen, once again indicating enhanced dominance of the deprived eye.

Two EEG study applied a **binocular frequency tagging** approach [34, 65], similar in all respects to the dichoptic frequency tagging described above but using congruent stimuli in the two eyes, that are readily fused despite their different flicker frequency. Results were reported as deprived/non-deprived eye response ratios (i.e. the ratios of the Fourier amplitudes at the harmonics of the corresponding stimuli, measured from occipital electrodes), which was shifted in favor of the manipulated eye following monocular deprivation [34] and a more complex monocular manipulation [65]. In a conceptually similar approach [36], both eyes are presented with a luminance patch of variable intensity, uncorrelated between eyes. Correlating the EEG signal from occipital electrodes to each temporal series allows for estimating the impulse response function for each eye. Its peak amplitude measures the contribution of each eye to the evoked response, which was enhanced for the deprived eye and suppressed for the non-deprived eye. Another parameter of the impulse response function is the oscillatory pattern observed after the initial peak; this echo response is dominated by an alpha rhythm, associated with perceptual sampling. In contrast with the modulation of visual evoked responses, the same dataset revealed no significant alpha modulation in the echo response – though the deprived eye amplitude was numerically larger after vs. before deprivation.

#### Monocular techniques

Monocular techniques measure sensitivity in the two eyes separately; these may be combined to compute an ocular dominance index, under the key assumption that enhanced monocular sensitivity implies enhanced dominance.

Studies measuring **monocular contrast sensitivity** provided partially conflicting results, potentially related to differences in the type of patch (translucent vs. light-tight). In three studies [2, 21, 60], application of a translucent patch was found to enhance sensitivity in the deprived eye (two of these also tested the non-deprived eye and found impaired sensitivity [2, 21]), while no modulation of deprived eye sensitivity was observed in Lunghi et al. [1]. Where the whole contrast-sensitivity function was measured, a preferential enhancement of the higher spatial frequencies was revealed [60] – but no sensitivity modulation was observed when stimuli were embedded in noise. Another four studies [25, 38, 43, 57] tested the effects of a light-tight monocular patch, which primarily impaired sensitivity in the non-deprived eye (seen in three out of four studies), with only a marginal enhancement of sensitivity in the deprived eye (only observed in [25]). Similar modulations were observed for a more complex monocular manipulation [25]. The one study that failed to report a sensitivity modulation [38] measured contrast-discrimination thresholds rather than absolute thresholds, i.e. participants' ability to report a contrast difference between two halves of a suprathreshold (clearly visible) grating. Another null finding came from **monocular global motion coherence** thresholds [25], which did not change following the application of an opaque patch (in contrast with the results of dichoptic global motion coherence) indicating that monocular sensitivity to motion is not affected by deprivation.

Few studies used EEG or fMRI to study the amplitude of responses evoked by monocular stimulation [8, 11, 23, 49, 58]; all achieved monocular deprivation through the application of a translucent patch. Lunghi et al. [8] used EEG to measure the **event related potentials** for individual stimulus presentations and found that deprivation modulates the amplitude of the earliest component of visual evoked potentials, known as C1. This was enhanced for the deprived eye and suppressed for the non-deprived eye. Federici et al. [58] replicated the C1 modulation; in addition, they quantified the spectral composition of EEG oscillations induced by stimulation. These consist of oscillations that are neither time- nor phase-locked to the stimulus onset, yet they are induced by the stimulation and modulated by its content. After monocular deprivation, alpha oscillations induced by deprived eye stimulation were reduced. Assuming that induced alpha is associated with inhibition, the modulation was interpreted as enhanced excitability, consistent with the modulations of evoked responses. Zhou et al. [11] measured **steady-state evoked potentials**, extracting the Fourier amplitude of the occipital EEG signal at the first harmonic of the monocular stimulus frequency. The results showed a selective enhancement of the deprived eye response, with no corresponding suppression of the non-deprived eye. Using ultra-high field functional MRI, Binda et al. [23] measured **evoked BOLD responses** in the primary visual cortex and Kurzawski et al. [49] extended the analyses to the visual thalamus. The results showed enhanced deprived eye responses and suppressed non-deprived eye responses in the primary visual cortex, particularly for the higher spatial frequencies tested, but no modulation in the lateral geniculate nucleus, supporting a cortical origin of the effect. However, a reliable modulation of evoked responses was observed in the ventral pulvinar.
